# JBrowse Connect: A server API to connect JBrowse instances and users

**DOI:** 10.1101/699504

**Authors:** Eric Yao, Robert Buels, Taner Z. Sen, Lincoln Stein, Ian Holmes

## Abstract

We describe JBrowse Connect, an optional expansion to JBrowse that allows live messaging via WebSockets, notifications for new annotation tracks, heavy-duty analyses initiated by the user from within the browser, and other dynamic features. We present JBlast, an example application of JBrowse Connect that allows users to specify and execute BLAST searches, managed by a Galaxy instance, as well as tracking job progress and viewing results, all in the context of the browser.

## Introduction

JBrowse (Buels et al. 2016) is a dynamic HTML genome browser installed at over 3,000 sites with over 40,000 monthly active users (Google Analytics, May 2019) and a thriving ecosystem of over 50 plugins, many contributed by third parties unaffiliated directly with the project (https://gmod.github.io/jbrowse-registry/). As one of the first JavaScript client genome browsers with a web API that can work directly from a static set of files, JBrowse can be run using any web server, including very lightweight web servers.

By design, the communication of information in JBrowse is one-way; there is no concept of a session or persistent connection, and no way for clients to push messages back to the server, or to other peer clients. For many purposes, this static model can be advantageous. For example, it is easy to deploy a genome as a “static site” (a set of files that are simply served up with no processing by a web server, and so can reside in cheap storage, e.g. in an Amazon S3 bucket). The static model is also inherently secure, as no code needs to be run on the server after the initial deployment. Thus, for many applications, the a client-only genome browser is ideal. However, for some specialized purposes - such as if the genome browser is intended to be not just as a vehicle for displaying information, but a portal to initiate analysis - it can be limiting that the client and server have no way to talk to each other during a browsing session.

As a partial workaround for this limitation, many groups have integrated some kind of server-side processing into JBrowse; either by having it be the last step of a workflow that generates and indexes static files, or by integrating it more deeply into a content management system. Some of these integrations are re-usable in general contexts. Notably, the Apollo project enables biocurators to collaborate in real time editing genome annotations (Lee et al. 2013; Dunn et al. 2019). Bioinformatics web applications such as Tripal (Ficklin et al. 2011) and Galaxy (Afgan et al. 2016) have extensions that integrate JBrowse to some extent. JBrowse’s plugin framework facilitates this sort of customization. However, since JBrowse was originally designed as a client-side application, with no API for dynamic server integration, there are limits to how smoothly such extensions can be integrated with the JBrowse user interface.

In this paper, we present JBrowse Connect, an optional and generic server extension to JBrowse that runs simultaneously as a JBrowse plugin (on the client) and as an extensible back-end (on the server). JBrowse Connect is designed to serve client requests for management and analysis of genomic data; to broker messaging between client and server (for example, to notify logged-in users of newly-added annotation tracks); to enable integration with 3rd-party analysis tools; and to provide mechanisms for straightforward application-specific extensions (“hooks”) on both client and server. It leverages Sails, a JavaScript model-view-controller framework with object-relational business logic and websocket integration. Like the JBrowse client, with its plugins, JBrowse Connect is extensible. As an illustration of how it can be extended, we also present JBlast, a JBrowse Connect hook that bridges a JBrowse client and a running Galaxy instance (Goecks et al. 2013; Afgan et al. 2016). As an example analysis task, we have implemented BLAST search. To users, this example is intended to demonstrate the experience of initiating a nontrivial analysis from within JBrowse; to developers, it demonstrates the API for developing a JBrowse plugin that triggers, monitors, and then collects the results of an analysis workflow in Galaxy (or potentially another workflow manager).

JBrowse Connect is an entirely optional extension to JBrowse. For site administrators who prefer the strong security and light CPU footprint of static hosting, JBrowse is (and will remain) usable without the extra server component. Further, JBrowse Connect is backward-compatible: if a JBrowse instance is in place, but evolving requirements have revealed a need for a more robust server component (whether it be to notify users of new tracks in real time, to allow them to initiate analyses or workflows from the browser, or for other reasons), then JBrowse Connect can be configured to work with an existing JBrowse instance with minimal disruption: JBrowse Connect uses the same filesystem structure and file formats as JBrowse, so that indexing or data files do not need to be regenerated or modified in order to install the server component.

In the Results section, we narrate the user’s experience of initiating and inspecting the results of a BLAST search from within JBrowse, using the example JBlast plugin. Then, in the Methods section, we outline the technical architecture of JBConnect (the core server software for JBrowse Connect) and JBClient (the corresponding JBrowse client plugin).

## Results

A JBrowse instance with the JBlast plugin installed looks mostly identical to vanilla JBrowse, but new buttons appear in the user interface. Using these buttons, the user can select a region of the reference genome from within the genome browser and, with a single mouse click, dispatch it for search against an instance-specific BLAST database. The user may alternatively submit a query sequence and BLAST it against the reference genome (these two alternative models are referred to as “reference-as-query” and “reference-as-target”). The progress of the BLAST search may be tracked through a popup queue window from within the user’s JBrowse instance. When the results are available, they show up immediately in the user’s web browser as a new track. A new client plugin allows the results to be dynamically filtered.

The individual steps in this process are illustrated using a flowchart in Figure 1, and using screenshots of the genome browser in Figure 2 and Figure 3.

**Figure 1:**
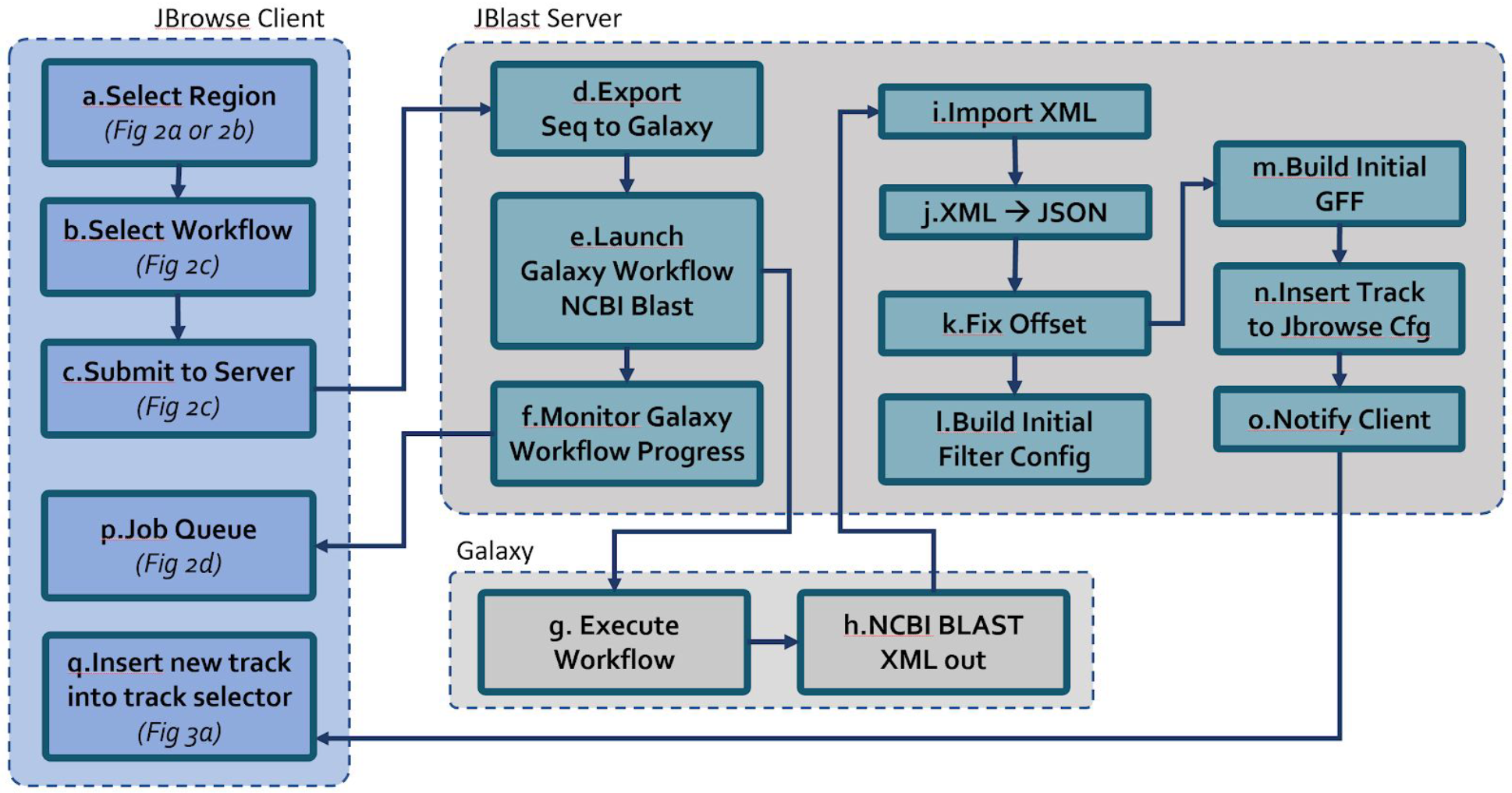
JBlast analysis involves coordination between the JBrowse client and a Galaxy instance with a BLAST database, mediated by the JBrowse server. The user first initiates a BLAST query from JBrowse (a,b). Clicking submit c) uploads any required parameters (like the query sequence) and queues the job for execution. The job queue, having received the job, exports the query sequence to Galaxy (d) and launches the corresponding Galaxy workflow (e). Throughout the job execution cycle, JBConnect monitors the progress of the job and workflow (f) and updates the client’s Job Queue panel (p). Galaxy executes the workflow (g) and upon completion produces a resulting BLAST XML file (h). Then, JBlast performs the post processing steps i through o, preparing the result for JBrowse rendering. The job completes as the resultant track appears in JBrowse (q). Once the job is submitted, it will complete regardless of whether the JBrowse client is running or not.

**Figure 2:**
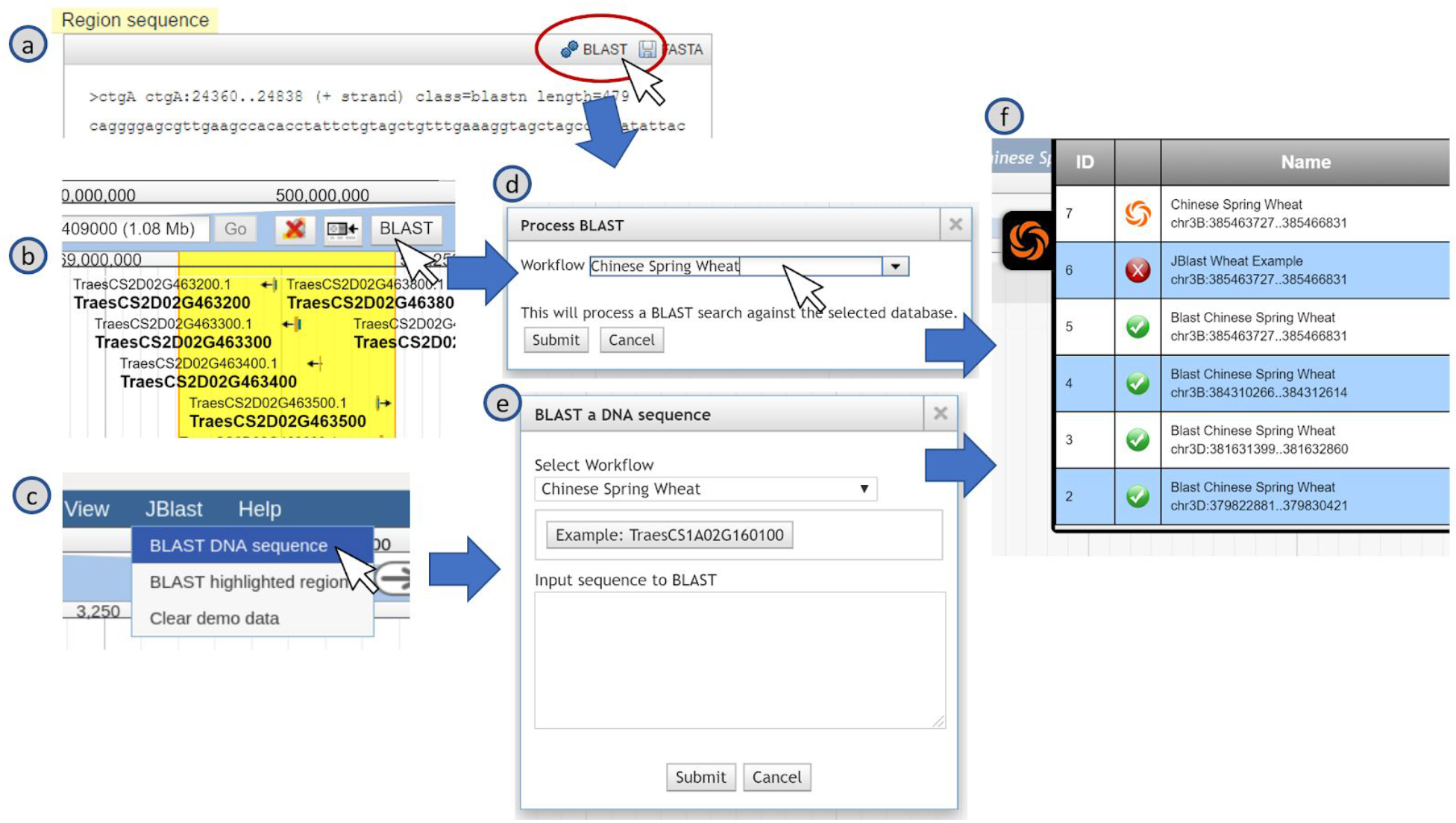
JBlast offers three ways to submit a BLAST query: a) select an existing genome annotation feature, b) highlight a region, or c) manually enter a sequence. The first two methods (a,b) use the “reference-as-query” model: the user selects a piece of the reference genome as a query, and the administrator configures an independent database of sequences to be the targets. In contrast, the third method (c) uses the “reference-as-target” model: the user supplies their own query sequence and the reference genome is configured as the target. Each method brings up an appropriate dialog (d,e) from which clicking the Submit button will launch the Job and open the Job Queue panel (f). The JBrowse instance with the JBlast plug-in is shown here based on the tracks generated for the Chinese Spring IWGSC RefSeq v1.0 wheat genome assembly (International Wheat Genome Sequencing Consortium (IWGSC) et al. 2018).

**Figure 3:**
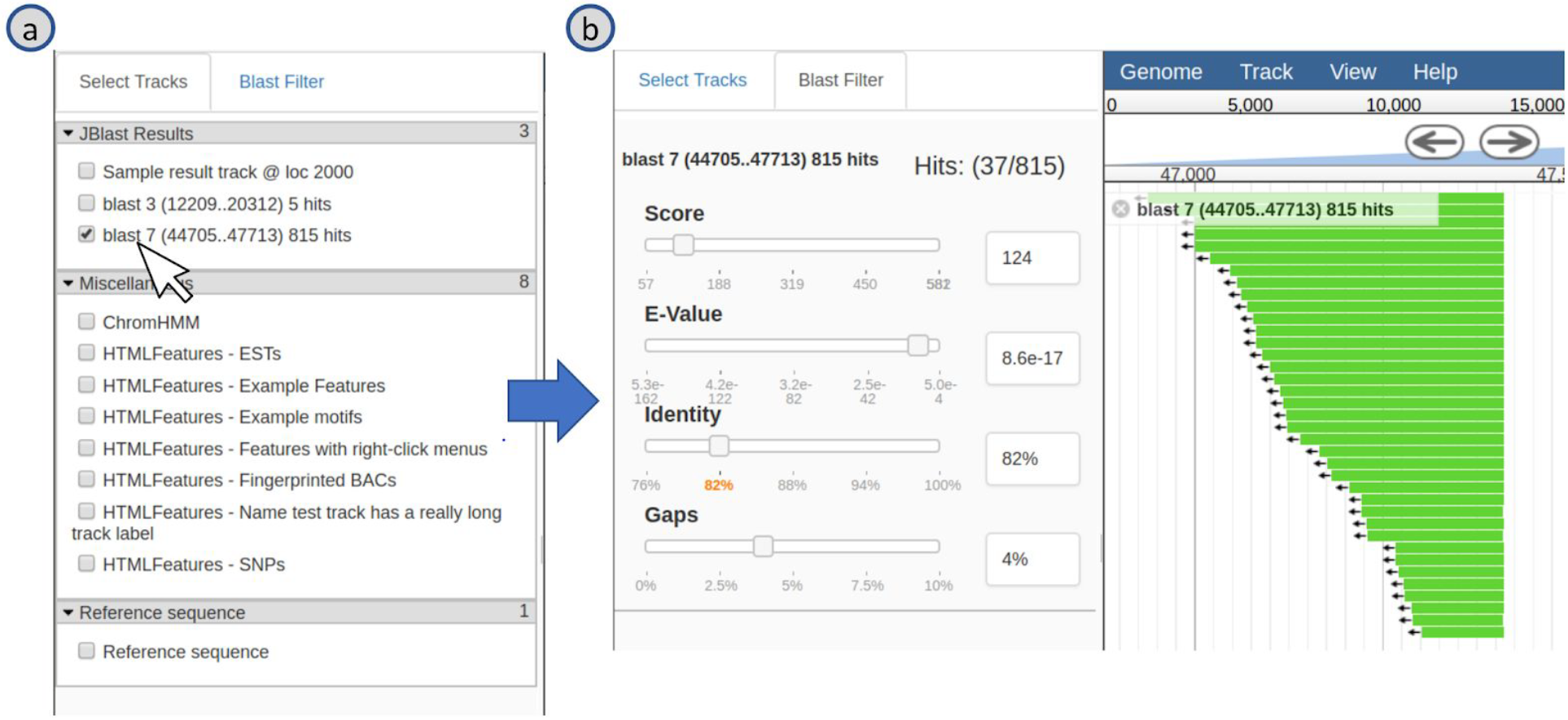
Analyzing results of a JBlast query (initiated in Figure 2). As soon as results are available, they appear as a new track in the user’s track selector (a). Once the user selects the track, it will appear in the main view, accompanied by a BLAST Filter panel (b) that allows the user to interactively and dynamically filter results by score, E-value, sequence identity, or alignment gappiness.

The JBlast plugin uses a Galaxy workflow to perform its BLAST analysis. Our original intention, developing this software, was to model all post-processing and format conversion steps as tasks within the Galaxy workflow, to facilitate system administration through the Galaxy interface. However, as we developed this system, it became apparent that requiring Galaxy as a non-optional dependency would significantly complicate installation in many cases, and would likely narrow the potential user base. Further, contrary to our intention of fostering re-use by exposing tasks through Galaxy, integrating these tasks tightly with the Galaxy code made it harder to create simple stand-alone workflows that managed tasks in a different way (e.g. through a local job queue). Consequently, we factored some of these operations into the JBlast core (Figure 1).

As a result of this design, JBlast (and JBConnect) are not limited to Galaxy. In principle, JBrowse Connect plugins can talk to other back-end bioinformatics services, including scientific computation providers such as Cyverse. They can also readily just run native code on the host, or act as simple proxies for other queuing systems. Indeed, the JBlast project provides and out-of-box example of stand-alone BLAST example.

Figure 2 shows the three methods to submit a BLAST query using a real-world example. The tracks were taken from the GrainGenes Genome Browser page (https://wheat.pw.usda.gov/GG3/jbrowse_Chinese_Spring) for the Chinese Spring IWGSC RefSeq v1.0 wheat genome assembly (International Wheat Genome Sequencing Consortium (IWGSC) et al. 2018). The first method (2a) is to select an existing feature from an existing track. In the feature’s Details dialog box under the Region sequence a blast button available. The second method (2b) is to use JBrowse’s highlight feature. Upon highlighting a region, a Blast button will appear on the toolbar. The 2nd method is also accessible from the JBlast menu, as shown in 2c (“BLAST highlighted region”). Clicking the Blast button from method 2a and 2b will lead to Process BLAST dialog (2d) which allows the user to choose the Workflow that will execute the BLASTn operation. Upon submitting the job, a new job will appear in the Job Queue panel (2f).

A third method (2c) is accessed from the JBlast menu, “BLAST DNA Sequence.” Selecting this item leads to the Blast a DNA Sequence dialog, which allows for entering an arbitrary DNA sequence by pasting sequence (with Ctrl-V). Again, upon submitting the job, a new now will appear in the Job Queue panel (2f).

### Case Study: Discovering syntenic regions in a polyploid plant

In order to show the usefulness of JBlast in a real-world example, we will use the demo JBrowse instance with the JBlast plug-in for the Chinese Spring IWGSC (International Wheat Genome Sequencing Consortium) RefSeq v1.0 wheat genome assembly (International Wheat Genome Sequencing Consortium (IWGSC) et al. 2018). The demo JBrowse instance can be accessed at a GrainGenes development site at http://graingenes.org:1337/jbrowse/?data=IWGSC (Blake V.C., Woodhouse M.R., Lazo G.R., Odell S.G., Wight C.P., Tinker N.A., Wang Y., Gu YQ, Birkett C.L., Jannink J.-L., Matthews D.E., Hane D.L., Michel S.L., Yao E., Sen T.Z., “GrainGenes: Centralized Small Grain Resources and Digital Platform for Geneticists and Breeders,” Database: The Journal of Biological Databases and Curation, in press). As a simple security feature, users must authenticate; a guest login/password pair (juser/password) are provided for evaluation purposes.

In contrast to a BLAST session performed using command-line or on a customized BLAST interface, JBlast offers immediate integration of the resulting sequence alignments with JBrowse’s annotation visualization capabilities. JBlast results appear as tracks, allowing in-depth exploration of genomic elements and annotations proximal to the JBlast hits.

The Chinese Spring wheat genome provides a good case study. It is a hexaploid plant, with six sets of seven chromosomes (2n=6x=42). BLAST hits are expected to be distributed across homoeologous chromosomes, which are historically named as A, B, and D (Akhunov et al. 2003; Glover, Redestig, and Dessimoz 2016).

The example sequence provided in the dialog box for the “BLAST DNA sequence” is a truncated 5’ sequence of the TraesCS1A02G160100 gene, located on chromosome 1A, which codes for a ribosome-associated protein (according to the Gene Ontology annotations for the gene). When users choose the example as their query sequence, the BLAST job should generate four hits. Setting the sequence identity threshold in the BLAST hit filter to 90% should winnow this down to three hits, listed for convenience in the filter, and ordered according to the BLAST E-value of the hit. These hits are on chromosomes 1A, 1B, and 1D. As shown in Figure 4b, the BLAST alignment for a given hit can be displayed by clicking on the displayed feature corresponding to that hit. Then, by adjusting the zoom level, syntenic regions in the homoeologous chromosomes can be identified, examined, and analyzed for further study.

**Figure 4:**
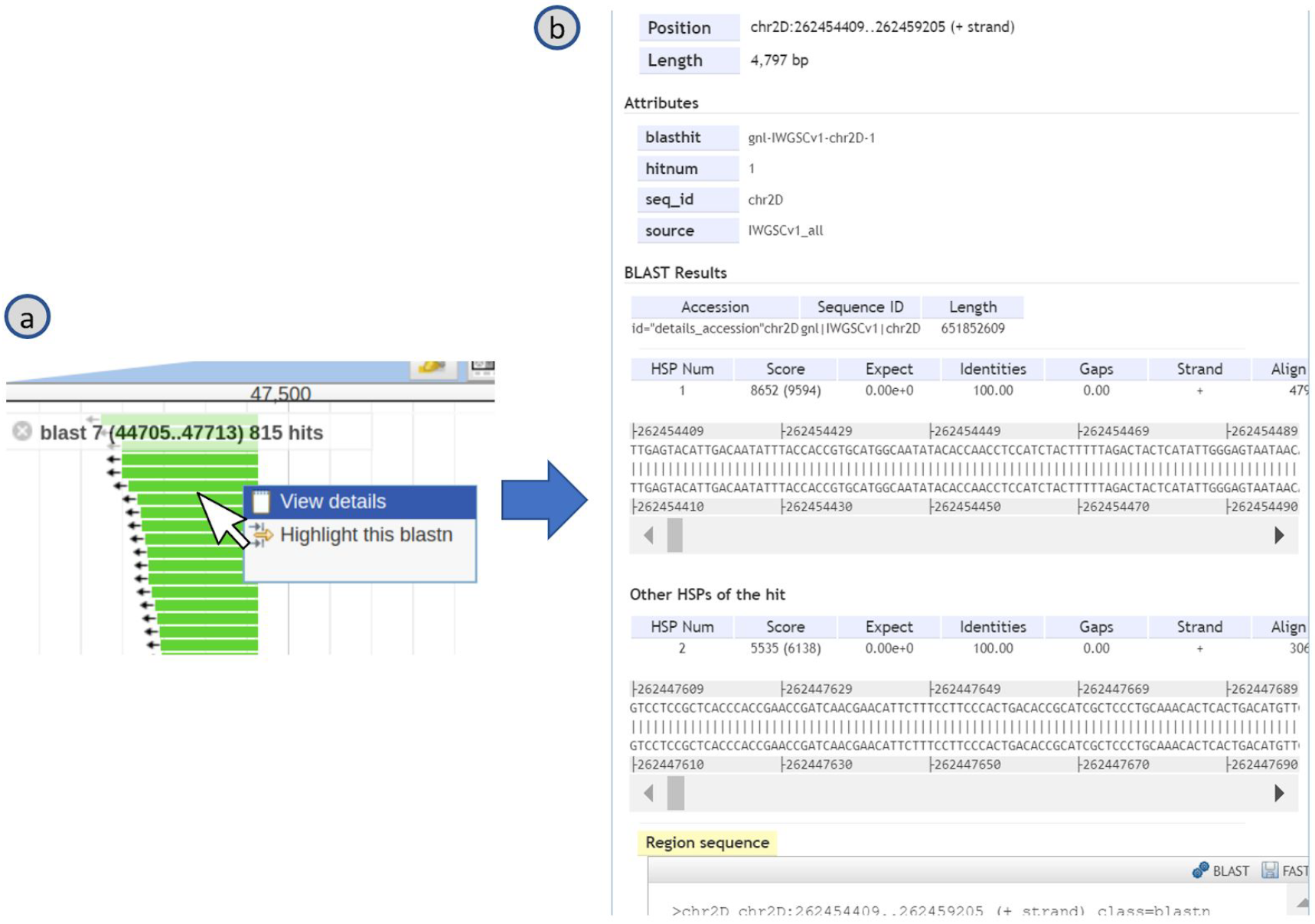
The features in a BLAST result track (a) are BLAST hits, the details of which can be viewed in the feature details dialog box (b) by double clicking the feature or access via its right-click menu. The High-Scoring Segment Pair (HSP) is provided for the given hit, and other HSPs from the same hit are also provided.

**Figure 5:**
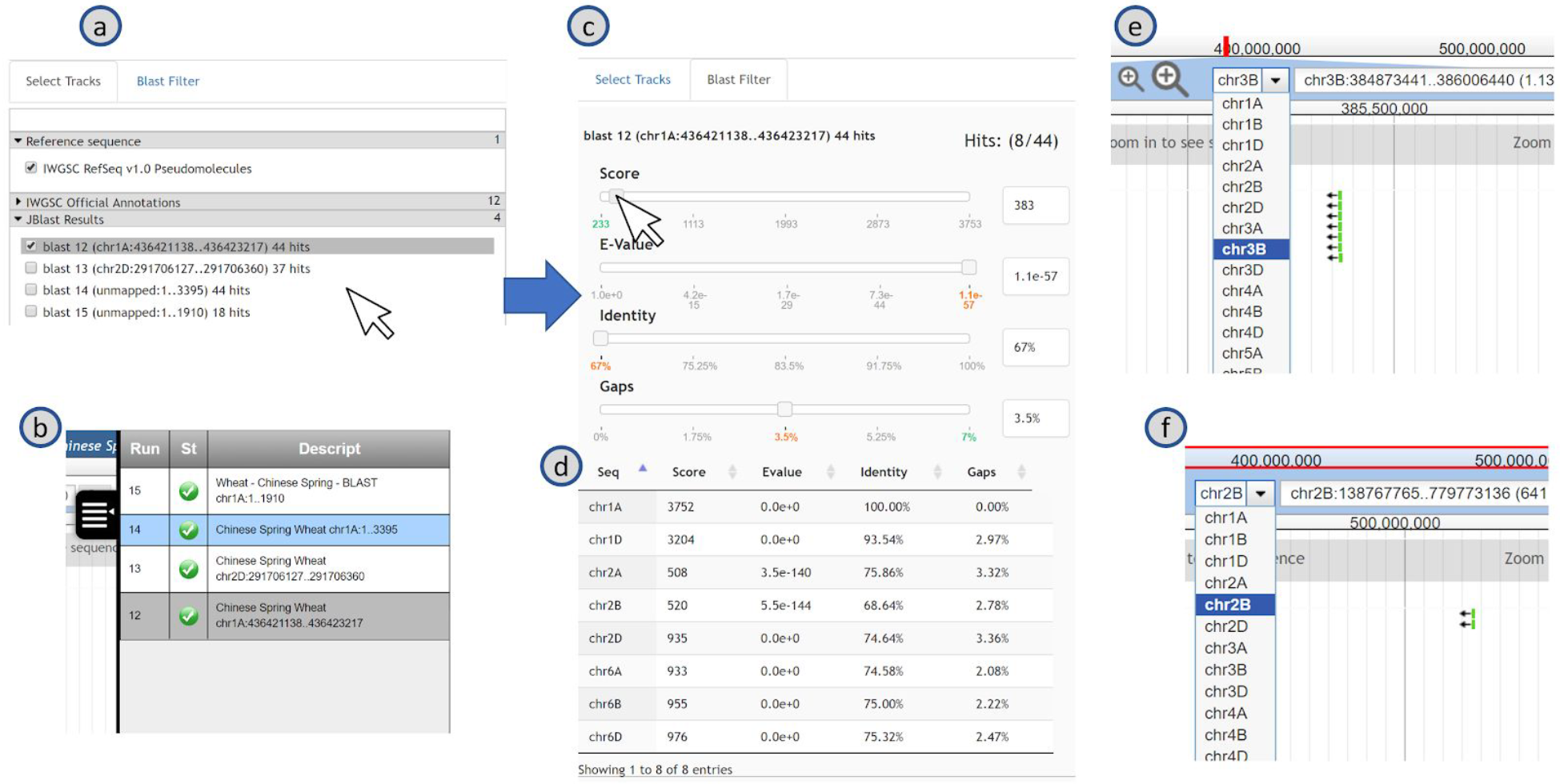
Analysis of the results of a BLAST search made using the online JBlast demo at GrainGenes, which is configured to use Chinese Spring Wheat as the reference genome. In the “reference-as-target” workflow used for this search, the BLAST database is identical to the reference genomes (i.e. the user contributes a sequence to be searched against the reference genome, in contrast with the other “reference-as-query” use case implemented by JBlast, where the user selects a region of the reference genome and searches it against an entirely separate BLAST database). In the reference-as-target model, BLAST hits can occur anywhere in the genome and so can be located on multiple different chromosomes. The result track can be selected from the track selector (a) or from the job queue (b). Once the user selects the track, it will appear in the main genome browser view, accompanied by a “BLAST Filter” panel which includes the filter sliders (c) that allows the user to dynamically filter results and the Result Table (d) which shows the filtered results and allows the user to click the row to jump to the selected location. Examples of two views in two chromosomes are shown in (e) and (f).

In this case, three hits are aligned with the following high-confidence gene models: TraesCS1A02G160100 (self) on Chromosome 1A (sequence identity: 100%), TraesCS1B02G176500 on Chromosome 1B (sequence identity: 93%), and TraesCS1D02G157300 on Chromosome 1D (sequence identity: 93%). According to the official annotations provided by the IWGSC, these three high-confidence gene models have the following common gene ontology (GO) annotations: GO:0005840 (ribosome), GO:0003735 (structural constituent of ribosome), GO:0008097 (5S rRNA binding), GO:0006412 (translation), GO:0008152 (metabolic process).

These results illustrate how JBlast enables quick identification of duplicate gene models, which can be further analyzed by comparing and studying the biological information embedded in other JBrowse tracks. This case study demonstrates not only the simplicity and power of using JBlast in studying polyploid species, but also provide evidence that users will be able to harness JBlast’s capabilities as easily to perform cross-species analyses.

## Methods

The operation of JBrowse Connect involves substituting a dynamic server program (JBConnect) for the static web server that normally serves up a JBrowse instance. JBConnect operates as a web server and service, presenting the usual JBrowse application as a route, along with the JBrowse client application (a set of JavaScript, CSS, and HTML files), and a number of other web services implemented as a RESTful API. Among other functions, JBConnect can inject additional plugins into the libraries that constitute the client; one such injected JBrowse plugin is JBClient (bundled with JBConnect), which augments JBrowse with a login panel and a Job Queue panel (Fig 2d), a clean and simple user interface for viewing queued, failed, completed, and active jobs. It illustrates a straightforward use of JBConnect’s queue as a simple interface to more powerful workflow engines such as Galaxy, which in turn connects to many kinds of cluster- and cloud-based workflow management systems.

### JBConnect Server Architecture

JBConnect is a Sails.js application. As such, it is built on node.js and Express, with a model-view-controller architecture, a built-in object-relational mapping with multiple back-end database handlers (Waterline ORM), a websockets API (Socket.io) with sub-pub support, a flexible authentication service (passports.js), and a framework called Blueprints which ties models, controllers, database objects into a consistent server API and REST interfaces between client and server, providing notification to clients of database updates, all around a highly scalable architecture. JBConnect is written in 100% JavaScript.

Figure 6 shows the JBConnect server architecture and how it extends the Sails architecture. Job Queue is an encapsulation of Kue, a Javascript priority queue management service. Each entry in the queue is represented by a Job object. Closely tied to the queue is the Job Service Interface, which is represented by a Service object. A job service may implement adapter interfaces to 3rd party server applications (local or remote), such as Galaxy. The Job Service may also implement REST interfaces specific to the service. The JBrowse Plugin Service implements a mechanism that allows one or more JBrowse plugins to be injected into JBrowse along with any dependent modules it may rely upon. Dataset/Track Services provides an encapsulation of JBrowse datasets and tracks. It maintains synchronization with JBrowse trackList.json configurations while also providing notification to clients when there are changes in track/dataset configurations. The Commands framework provides a mechanism for the command line tool, jbutil, to be functionally extended by JBConnect hooks. Configuration management provides a configuration service that aggregates configurations by the server and by JBConnect hooks, as well as the user-defined configurations specific to a particular setup.

**Figure 6:**
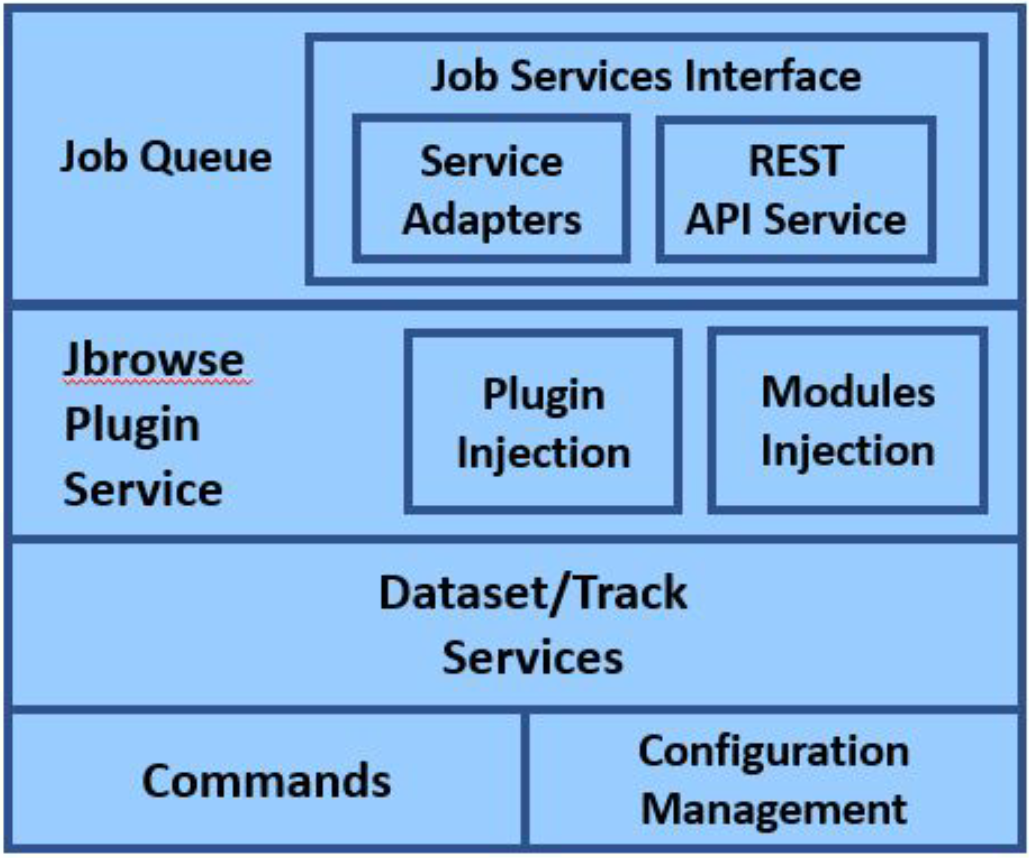
JBConnect Server Architecture

### Object-relational datasets and tracks

The core database tables of JBConnect, termed “model objects” in Sails’ object-relational mapping, represent the data structures required to perform asynchronous remote analysis in a genome browser.

Two of these model objects, “Dataset” and “Track”, represent core concepts of the JBrowse genome browser, as described previously (Buels et al. 2016). A Dataset (which typically corresponds to an individual genome) consists of a set of reference sequences and associated Tracks, which are annotations that may be displayed in the browser. When these entities are manipulated through JBConnect’s API, the static files that the JBrowse uses as indices (of tracks and datasets) are automatically updated, and notifications are sent to all connected JBrowse clients that the tracks (or datasets) have been changed and should be refreshed.

The new “Job” model object encapsulates analysis jobs. Using the JBConnect API, jobs may be submitted, listed, updated, or destroyed. The JBClient plugin uses this API to manage jobs. Two other new model objects, “Passport” and “User”, provide a flexible interface to local authentication mechanisms and user accounts.

### JBConnect Hook

JBConnect Hooks framework leverages Sails Installable Hooks framework and extends it in a number of ways, with the intention of providing a mechanism of expanding JBConnect’s functionality using an NPM module approach.

There follows a summary of the JBConnect Hook. A more detailed description can be found in the online documentation: https://jbconnect.readthedocs.io/en/latest/hooks.html

Figure 7 shows the JBConnect Hook framework that provides the following ways for expanding JBConnect:

- Job services are the main mechanism for processing a job. The job service may include an adapter for local or 3rd party server API support, such as Galaxy API. REST APIs specific to the service can also be part of the Job Service. The Job Queue relies on the job service to provide translated state information and execution pre and post processing of the analysis operations. The REST API service of the Job Service is an alternative to the standard sails controller REST interfaces.
- JBrowse plugin / injection (plugins that are tightly integrated with the server-side hooks) along with client-side module dependencies, used by the JBrowse plugins. The injection occurs upon sails lift and copies the necessary plugins into the JBrowse server plugins directory.
- Model/controllers/services as provided by Sails Blueprints. These services are merged with native server models/controllers/services into globally accessible objects.
- Command extensions - command options and implementation are merged with the function of JBConnect’s jbutil command, providing extended command capabilities specific to the JBConnect hook.
- Configurations are aggregated with the JBConnect server configurations.

**Figure 7:**
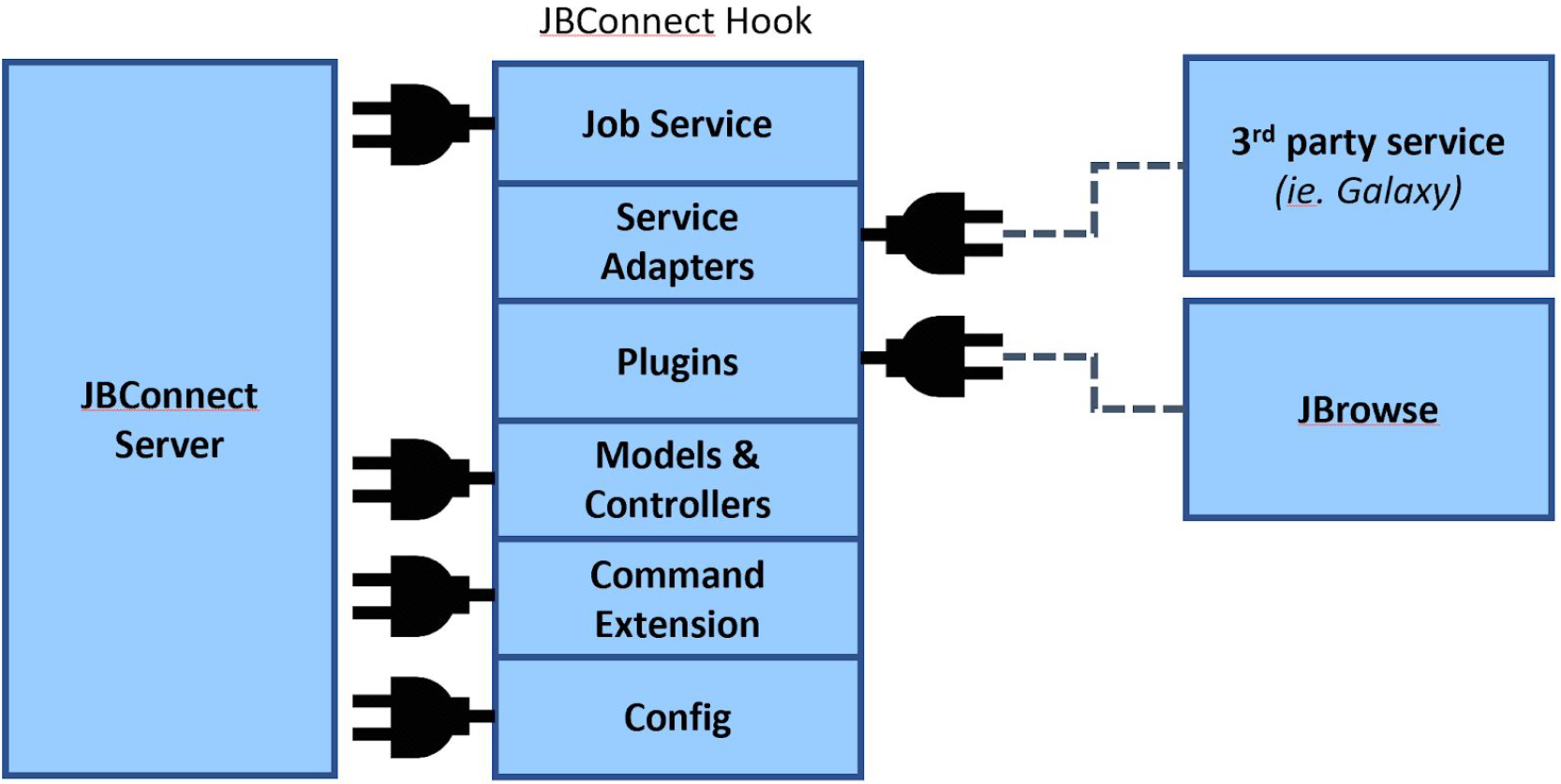
JBConnect Hook

A tutorial is available on creating a job service from scratch is provided, illustrating required methods for job execution: https://jbconnect.readthedocs.io/en/latest/tutorials.html

### Client plugin

JBClient is the client counterpart to the JBConnect server. It is a client-side plugin for JBrowse that extends the JBrowse graphical user interface and communicates with the JBConnect web services. The JBClient plugin is automatically injected by JBConnect when the server is started. It extends the JBrowse user interface with the login panel and the job queue panel (Fig 2d), and automatically responds to new or modify track events by updating the associated JBrowse views.

## Discussion

JBrowse has previously been integrated with genomics-oriented content-management systems such as Tripal (Ficklin et al. 2011), as well as computation services such as Cyverse (Merchant et al. 2016) and bioinformatics pipeline managers such as Maker (Holt and Yandell 2011), SeqWare (O’Connor, Merriman, and Nelson 2010), and Galaxy (Goecks et al. 2013). JBrowse has been implemented as a tool in Galaxy (https://biostar.usegalaxy.org/p/15491/). Galaxy also has its own genome browser, with a streamlined set of features compared to JBrowse, from which it is possible to launch and track Galaxy jobs. A basic “BLAST” button (lacking Galaxy integration or queue monitoring) has been available for some time to biocurators using the genome annotation editor Apollo (Lee et al. 2013), which is built using JBrowse.

JBrowse Connect builds on these previous efforts to offer a general-purpose framework for doing analysis from within the established, richly-functional, and well-supported JBrowse genome browser. The user experience is well-integrated with browsing: the user can maintain focus on biological objects of interest (genomes and genome annotations) without making unnecessary detours to a separate workflow-management application. The analysis queue is a minimally-invasive popup window accessible from within JBrowse. BLAST results appear immediately within their genomic context, and are instantaneously available for other users of the site to share. This can all be handled in a manner complementary with the local policies and server software for authentication, access control, and content management.

Aside from performing alignments or sequence homology searches, the JBrowse Connect infrastructure can in principle be used for many common tasks around genomics web development (and content management more generally) such as persistent user uploads (and data-sharing), automatic indexing of uploaded files, exposure of more JBrowse functions as server-hosted web services, and policy-based access controls.

As bioinformatics web applications develop, genomes (and genome annotations) are arguably one of the most materially intuitive data structures we have. A transcript is linear not just because it represents a conceptual ordering that is isomorphic to a linear graph, but because it represents a real-world object that has the topology of a line. Thus, it represents a natural “social object” (Engeström 2005). In this context, it seems likely that more and more applications will coalesce around the genome browser, either as a chassis or as a common component. The JBrowse Connect system aims to accelerate this process by implementing many of the common tasks required to turn a modern static web experience into a dynamic, collaborative one.

## Appendix: JBlast Hook

### Overview

JBlast (jblast-jbconnect-hook) is a JBConnect hook module. It provides server-side integration with both JBConnect and JBrowse that enables Blast analysis and is tightly integrated with JBrowse. JBlast can launch stand-alone NCBI blast jobs directly, or it can be configured to run blast through Galaxy, as a workflow. Through the JBrowse user interface the user can choose to submit an existing feature as a blast query, or to highlight a region to blast (Fig 2a, 2b, and 2c). The user can monitor the progress of the blast job through JBConnect’s job queue (Fig 2d) and the blast search results will appear directly as an inserted track in the track selector (Fig 3a). The resulting hits can be filtered through a Filter Panel (Fig 3b). The details of the hit can further be displayed in the Feature Details dialog box (Fig 4a).

### Job Services

JBlast implements two Blast analysis paths. One is a stand-alone method (basicWorkflowService) and another utilizes Galaxy (galaxyService).

basicWorkflowService is the stand-alone blast job service (Fig 7) that manages the lifecycle of a “local” NCBI BLAST job. The Job Service launches this job runner into action where it executes an NCBI blast command on the query data. The Job runner periodically monitors the state of the blast process to completion, while also updating the job processing engine on the status changes. Upon completion, it does the following:

- XML → JSON (Fig 7j) - Convert XML file into a JSON database as an associative array where the key is <Hit ID>:<HSP_num>, facilitating ease of addressability for future operations.
- Fix Offset (Fig 7k) - Fixes the offsets of the of the resultant hit features in the JSON database so that the resultant data can be presented in the domain of the original search region for the given reference sequence.
- Build Initial Filter Config (Fig 7l) - build the initial filter configuration file where filter settings are persisted for the resultant track asset.
- Build Initial GFF (Fig 7m) - based on the filter configuration file, the track presentation data is prepared in the form of a GFF file.
- Insert Track to JBrowse Cfg (Fig 7n) - The new track is inserted into the dataset configuration via the Track object. This essentially also pushes the data into the JBrowse trackList.json configuration file.
- Notify the Client (Fig 7o)- As part of the Track object insertion, the client (and indeed any listening client) is notified of the track insertion event Insert

**Figure 7:**
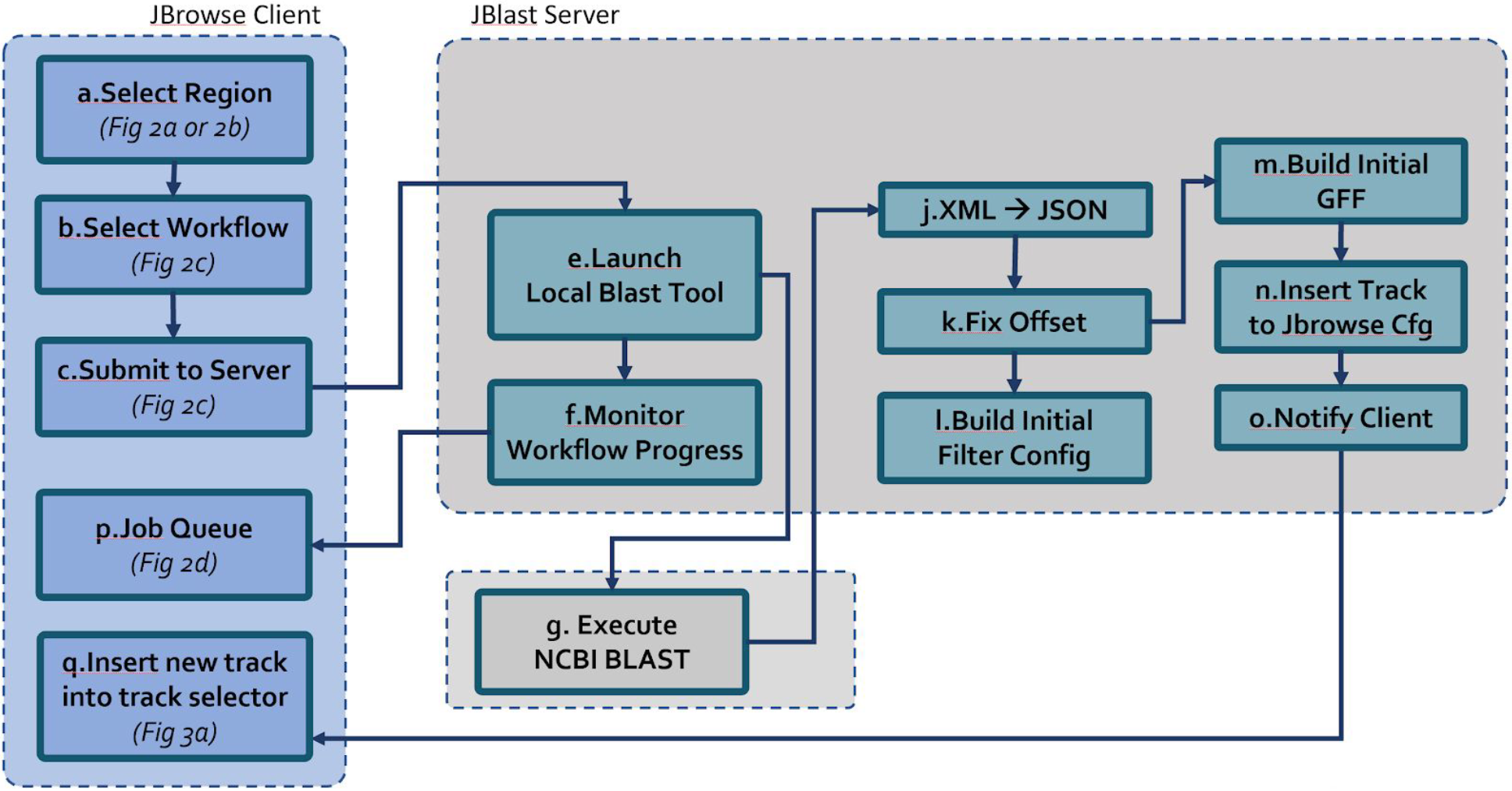
JBlast Stand-Alone BLAST analysis process. Similar to the Galaxy process (Fig 1), the user initiates a BLAST query from within JBrowse (a,b). Clicking submit (c) uploads any relevant parameters (such as the query sequence), and queues the job for execution. The job queue, having received the job, launches the corresponding local workflow script (e). Throughout the job execution cycle, JBConnect monitors the progress of the job and workflow (f) and updates the client’s Job Queue panel (p). The script begins processing BLAST (g) and upon completion produces a resulting BLAST XML file. Then, JBlast performs the post processing steps j through o, preparing the result for JBrowse rendering. The job completes as the resultant track appears in JBrowse track selector (q).

After completing the above stages, the job is considered complete.

Use of basicWorkflowService requires installing NCBI tools as well as configuring a blast databases. JBlast installs the git project enuggetry/ncbi-blast-tools which makes utilities available to automatically install the NCBI blast (utils/blast_getBlastUtils.js) and install NCBI blast databases (utils/blast_downloadDb.js). Another tool provided by the project (utils/blast_setPathDb.js) helps configure any arbitrary blast database for use by JBlast.

Local blast - configuring blast command line - blast profile

basicWorkflow supports a configuration option called BLAST Profiles. Blast commands have multi-faceted options that can be passed either through config file or directly in the job submission API.

The Galaxy adapter, galaxyService, is a job service that manages the lifecycle of BLAST that utilizes a Galaxy workflow in the execution of the job (See Fig 1), where the resultants are BLAST XML files. The Job Service launches this job runner into action where it uploads the query sequence to Galaxy and directs Galaxy to execute a BLAST workflow and the resultant is a BLAST XML file.

The Job runner periodically monitors the state of the Galaxy workflow to completion, while also updating the job processing engine on the status changes. Upon completion, it exports the resultant BLAST XML file from Galaxy and the proceed to perform steps 6.10 to 6.15 from figure 6 as described in basicWorkflowService, above.

Use of Galaxy requires setting up Galaxy with NCBI blast tools from the Galaxy toolshed and setting up a blast workflows that will export results in an XML format. This Job Service utilizes a adapter specific to communicating with Galaxy through the Galaxy API. The configuration of Galaxy API usage requires acquisition of an API key can be generated from the Galaxy user interface.

Job Services are a polymorphic interface where job runner services are an optional subclassed function of the Job service. JBlast’s galaxyService and basicWorkflowService from an API perspective and are functionally opaque to the client.

The filter management service is used by JBClient to apply filter changes to a particular BLAST result track asset. The client filter panel (Figure 3b) uses the API to change the filter thresholds for Score, E-Value, Identity and Gaps. GalaxyService and basicWorkflowService also use this to set the initial filter state, determining the max and min range of each result value.

## Acknowledgements

This work has benefited greatly from conversations with Nathan Dunn, Colin Diesh, Helena Rasche, Chris Mungall, Suzi Lewis, Deepak Unni, Alexie Papanicolau, Moni Munoz-Torres, and the GrainGenes team members.

## Funding

EY, LS, and IH were supported by NIH-NHGRI grant HG004483. RB, LS and IH were supported by NIH-NCI grant CA220441.

